# HMVA-DDI: A Hierarchical Multi-View Attention Network for Drug-Drug Interaction and Severity Prediction

**DOI:** 10.64898/2026.09.25.754374

**Authors:** Bibekananda Hati, Ruchika Bhat, Zakeerhussain D Kudari, Virendra Singh Kaira, P Sai Shankar, G. Jaiganesh

## Abstract

**Background and Objective:** Prescribing multiple drugs simultaneously, known as polypharmacy, is increasingly common in managing chronic diseases such as diabetes and cardiovascular conditions. Studies show that roughly 40% of elderly patients are on five or more medications, putting them at significant risk of adverse drug-drug interactions (DDIs) [1]. While computational methods based on Graph Neural Networks (GNNs) have shown strong potential for predicting these interactions from a drug’s molecular structure, most existing models are limited in two important ways: they rely on attention mechanisms that cannot adapt to the unique context of each atom in a molecule, and they only predict *whether* an interaction exists rather than *how dangerous* it might be.

**Methods:** We propose HMVA-DDI, a two-stage deep learning framework that addresses both shortcomings. In the first stage, our model predicts whether two drugs will interact. If it detects an interaction, the second stage classifies it into one of three clinical severity levels: minor, moderate, or major. To better capture molecular structure, we replace the standard attention mechanism with a more powerful version called Graph Attention Network v2 (GATv2), which considers each atom’s specific context when learning its representation. We also introduce a technique to prevent information loss in deeper network layers, and we tailor the input features separately for each prediction task.

**Results:** Evaluated on two standard benchmarks, DrugBank 5.0.3 [2] and DDInter [3], HMVA-DDI achieves an AUROC of 0.9819 and AUPRC of 0.9759 on binary interaction detection, a marked improvement over the baseline. For severity classification, our best model variant (qcSvr4) reaches an overall AUROC of 0.9444, with a particularly large improvement in detecting low-severity (Minor) interactions, a historically difficult class.

**Conclusion:** HMVA-DDI bridges the gap between computational drug research and clinical usefulness by not only predicting interactions accurately but also indicating interaction severity. This two-level system can help healthcare professionals prioritize warnings, reducing the “alert fatigue” problem where doctors are overwhelmed by too many non-critical notifications.

## 1 Introduction

Polypharmacy, defined as the concurrent use of five or more medications, has become an unavoidable reality in modern clinical practice. Driven by the aging global population and the rising prevalence of complex, long-term conditions, which include cardiovascular disease, type-2 diabetes, chronic kidney disease, and cancer, more patients than ever are managing multiple medical conditions simultaneously [4]. A 2023 meta-analysis found that approximately 39-40% of adults aged 60 and above are on five or more medications at any given time, with rates reaching as high as 80% in nursing home settings [1]. While this concurrent therapy is medically necessary, it exponentially raises the risk of adverse drug-drug interactions (DDIs), where one drug alters the behavior of another inside the human body. DDIs operate through two main pathways: *pharmacokinetic* interactions, where one drug changes how the other is absorbed, broken down, or excreted; and *pharmacodynamic* interactions, where two drugs amplify or cancel each other’s intended effects. These interactions can cause outcomes as mild as a slight reduction in a drug’s effectiveness, or as severe as organ failure, life-threatening bleeding, or cardiac arrest. Ensuring timely, reliable detection of DDIs is therefore not just a scientific challenge but a critical factor in patient safety and survival.

Historically, identifying DDIs has relied on exhaustive *in vitro* laboratory experiments and late-stage clinical trials. While these traditional empirical methods are indispensable for establishing ground-truth biological validation, they suffer from severe scalability bottlenecks. The sheer combinatorial explosion of possible drug pairs, including newly synthesized molecular entities and legacy drugs, makes it prohibitively expensive and temporally impossible to screen all combinations in a wet lab. This pressing bottleneck has driven the development of *in silico* computational methods designed to serve as an initial high-throughput screening mechanism to prioritize high-risk drug combinations for further biological testing. Early computational approaches largely relied on similarity-based heuristics, operating on the “guilt-by-association” premise [5]. These methods utilized matrix factorization [6] and classical machine learning over side-effect profiles or genomic targets. Although these foundational models paved the way for computational pharmacology, they relied heavily on hand-crafted features. They often failed to capture the highly non-linear, structural relationships that truly govern molecular interactions.

The advent of deep learning marked a massive paradigm shift in the field. Initial architectures, such as DeepDDI [7], leveraged one-dimensional SMILES (Simplified Molecular-Input Line-Entry System) strings to predict interactions. However, representing a complex three-dimensional biochemical entity as a flat, 1D string intrinsically destroys its rich topological and geometric information. To address this, researchers shifted to Graph Neural Networks (GNNs), which have become the state-of-the-art framework in this domain. GNNs operate directly on the 2D molecular graph, where atoms are represented as nodes and chemical bonds as edges. Landmark models like SSI-DDI [8] and its dual-view successor, DSN-DDI [9], showed that learning from both intra-molecular substructures (like specific functional groups) and inter-molecular dependencies yields superior predictive performance.

Despite the undeniable progress brought by GNNs, existing architectures, including the highly effective DSN-DDI, suffer from four distinct limitations that restrict their ultimate clinical utility: First, they predominantly rely on standard Graph Attention Networks (GATs). Standard GATs compute attention statically, meaning the relative importance of neighboring atoms is unconditioned on the query node, severely limiting the network’s expressive power when modeling complex, asymmetric biological bonds. Second, as molecular GNNs become deeper to capture global topology, they frequently fall victim to “over-smoothing”. This is a phenomenon where repeated message passing causes node embeddings to converge to indistinguishable values, effectively “squashing” the unique, low-level structural identity of the functional groups that actually cause the DDI. Third, previous models often use a rigid, uniform feature-engineering strategy. However, predicting whether an interaction exists is fundamentally different from predicting its clinical severity; the former benefits from global molecular descriptors, while the latter requires hyper-focused atomic-level precision. Finally, the vast majority of existing literature frames DDI prediction strictly as a binary task (Interaction vs. No Interaction). In real-world clinical decision support systems, binary alerts lead directly to “alert fatigue”, where physicians simply ignore warnings because the system flags every minor, clinically irrelevant interaction.

To address these challenges, we propose the **Hierarchical Multi-View Attention Network (HMVA-DDI)**, a novel two-stage deep learning framework engineered for both binary DDI prediction and multi-class severity classification. Our specific scientific contributions are as follows:

1. **Dynamic Attention Integration:** We replace the standard, static GAT with the strictly more expressive **Graph Attention Network v2 (GATv2)**. This enables dynamic, context-aware attention that correctly models the asymmetric dependencies between interacting molecular substructures.
2. **Over-Smoothing Mitigation via MVA:** We introduce a novel Multi-View Attention (MVA) module that acts as a robust **long-range skip connection**. By explicitly fusing feature representations learned at varying network depths, MVA prevents information squashing and allows the final prediction head to utilize both low-level atomic features and high-level global topology.
3. **Task-Specific Feature Augmentation:** We implement a bifurcated feature strategy. We augment atomic node features with 200-dimensional RDKit2D molecular fingerprints (capturing global physicochemical properties like LogP and polar surface area) exclusively for the binary interaction task, while reserving pure, unadulterated atomic features for the delicate severity classification task.
4. **Clinical Severity Stratification:** We hierarchically connect a binary interaction predictor to a multi-class severity classifier (Major, Moderate, Minor). By strictly conditioning the severity prediction on the existence of an interaction, we align the computational model with real-world clinical triage, directly combating alert fatigue.

Through these targeted innovations, HMVA-DDI not only achieves state-of-the-art predictive accuracy on benchmark datasets but also delivers the granular severity stratification required for genuine clinical deployment.

## 2 Literature Review

The field of computational drug-drug interaction prediction has gone through several distinct phases of development, each driven by new ideas about how best to represent drugs and their interactions mathematically. This section traces that journey from simple similarity scoring to today’s graph-based deep learning methods and critically identifies the gaps that motivate the design of HMVA-DDI.

### 2.1 Why Predicting DDIs Is Hard: The Biological Background

Before reviewing computational methods, it is important to understand what makes DDI prediction biologically difficult. Most drug interactions occur for one of two reasons.

First, two drugs may compete for the same biological machinery in the body. The liver is responsible for breaking down most medications, and a family of enzymes called Cytochrome P450 (CYP450) does most of this work. When two drugs both rely on the same CYP450 enzyme for elimination, one drug can block the other from being processed. The result is that the blocked drug builds up to dangerously high concentrations in the bloodstream. For example, if a patient takes warfarin (a blood thinner) and fluconazole (an antifungal), fluconazole blocks the CYP2C9 enzyme that breaks down warfarin, leading to warfarin accumulation and a risk of serious bleeding. Recent research has proposed integrating knowledge about CYP450 enzyme substrates directly into DDI prediction models to make them more biologically grounded [10].

Second, two drugs may act on the same receptor or biological target, either amplifying each other’s effects to dangerous levels (additive/synergistic toxicity) or canceling each other out (antagonism). For example, combining two drugs that both lower blood pressure can cause it to drop dangerously, even if each drug individually is safe at its prescribed dose.

This biological complexity means that predicting DDIs requires more than knowing the chemical formula of a drug. A model needs to understand whether two drugs share metabolic pathways, compete for the same protein targets, or have overlapping physiological effects. This context is crucial for understanding why certain machine learning approaches succeed while others fall short.

### 2.2 Early Computational Approaches: Similarity and Matrix Methods

The first computational approaches to DDI prediction were based on a simple but powerful idea: drugs that look and behave alike. If two pairs of drugs share a similar chemical structure, similar side-effect profiles, or similar protein targets, they are likely to have similar interaction patterns. This principle, sometimes called “guilt-by-association,” underpins methods like NDD [5], which built networks of drug similarities and used graph-theoretic techniques to propagate known interaction information across the network.

Matrix factorization methods, such as DDINMF [6], took a different angle inspired by recommender systems. Just as a streaming platform predicts what movies a user might like based on other users’ preferences, these methods factorize the known DDI matrix into latent representations that can fill in missing entries. The IPF method [4] extended this by integrating multiple similarity sources simultaneously.

Although effective for drugs with an extensive history in the database, these approaches share a fundamental weakness: the cold-start problem. When an entirely new drug enters the market with no prior similarity records or observed interaction history, using these models will produce unreliable predictions. This is especially critical for patient safety, since newly approved drugs carry the greatest risk of undiscovered interactions.

### 2.3 The Rise of Deep Learning on SMILES Strings

The arrival of deep learning brought new possibilities for representing drug structure. SMILES strings represent a drug’s molecular structure as a sequence of letters and symbols, much like a sentence describes an idea. DeepDDI [7] was the first major model to learn directly from SMILES strings, using a deep neural network to predict 86 different categories of drug interactions. Its success demonstrated that neural networks could extract meaningful structural information from chemical representations.

However, SMILES has a fundamental flaw for structural modeling: the same molecule can be written in many different ways depending on which atom you start with. Two drugs with nearly identical three-dimensional shapes might have very different SMILES strings, and vice versa. This ambiguity makes it difficult for sequence-based models to reliably learn structure-function relationships. Furthermore, SMILES strings cannot easily capture the three-dimensional spatial arrangement of atoms, the direction of functional groups, or the orientation of ring structures, all of which can be chemically decisive.

### 2.4 Graph Neural Networks: Learning from Molecular Structure Directly

The limitations of SMILES-based modeling naturally led researchers toward a more principled representation: the molecular graph. In a molecular graph, each atom is a node, and each bond is an edge. This representation is canonical where every molecule has exactly one graph, and it preserves all topological information about how atoms are connected. Graph Neural Networks (GNNs) were developed to learn from such graph-structured data by passing information between neighboring nodes in successive rounds of message passing.

DPDDI [11] was among the first to apply GCNs (Graph Convolutional Networks, a basic form of GNN) to DDI prediction. However, the most important advance came when researchers began modeling drug pairs simultaneously, rather than treating each drug in isolation. SSI-DDI [8] proposed that interactions between drugs are driven by specific functional groups, which are chemically active regions of the molecule, such as hydroxyl groups (−OH) or amino groups (−NH_2_). By building substructure-level attention into the GNN, SSI-DDI significantly improved accuracy.

Building directly on this, DSN-DDI [9] introduced a two-level attention system: one that attends to atoms within each drug (intra-graph), and one that attends to atoms across the two drugs being paired (inter-graph). This dual-view design let the model reason about how parts of Drug A might physically interact with parts of Drug B, rather than reasoning about each drug in isolation. DSN-DDI became the strongest baseline in the field and the direct predecessor to our work.

Other GNN architectures have explored complementary directions. DeepLGF [12] studied how best to combine local (substructure-level) and global (whole-molecule) features. GCNMK [13] used multiple graph kernels to capture different structural views of the same molecule. GraphSAGE-based adaptations [14] have been explored for settings with very large drug graphs where full neighborhood aggregation is computationally prohibitive. Attention-specific architectures like DRGATAN [15] and MHCADDI [16] demonstrated that the way attention is computed across drug pairs significantly affects the quality of learned representations.

### 2.5 A Critical Limitation: What Standard GATs Cannot Do

A key component used in most of the GNN models above is the Graph Attention Network (GAT), which allows each atom to weigh the importance of its neighbors before aggregating information. However, in 2021, Brody et al. [17] proved mathematically that the original GAT has a hidden limitation: it can only compute what they call “static” attention. This means the relative importance assigned to neighboring atoms does not depend on the atom that is asking the question. Imagine a student who always ranks the same book as most important, regardless of the assignment the ranking stays the same regardless of context. In chemistry, this is problematic because the importance of, say, a nearby oxygen atom depends entirely on the electronic environment of the atom it is interacting with.

GATv2 [17], the improved version, resolves this by computing “dynamic” attention where each atom’s view of its neighborhood is fully conditioned on its own current state. This seemingly small change has large practical consequences when modeling complex, context-sensitive bonds. Several recent works have shown that GATv2 improves molecular representation quality over standard GAT in property-prediction benchmarks. HMVA-DDI adopts GATv2 as its core message-passing primitive for exactly this reason.

### 2.6 Multi-Modal and Knowledge-Enriched Approaches

Researchers have increasingly recognized that molecular structure alone, however well-represented, cannot fully explain all DDIs. Many interactions are driven by biological mechanisms that are not visible in the drug’s chemical formula. For example, two drugs may not structurally resemble each other at all, yet both may inhibit the same CYP3A4 enzyme, making their combination dangerous.

This insight has motivated a wave of multi-modal models. GoGNN [18] incorporated Gene Ontology (GO) terms which are structured annotations describing the biological roles of proteins that gives the model context about what each drug does inside cells. DMFDDI [19] combined drug-target interaction data, enzyme information, and transporter data into a unified matrix factorization model. Knowledge-graph-enhanced approaches [10, 20] have begun embedding drugs into rich biomedical knowledge graphs where relationships between drugs, proteins, diseases, and pathways are all represented as interconnected nodes.

On the representation side, research on molecular fingerprints shows that different encoding methods capture complementary information. Extended Connectivity Fingerprints (ECFP) excel at capturing local, topological features around each atom. RDKit2D descriptors, by contrast, summarize global physicochemical properties such as how fat-soluble a drug is, how easily it dissolves in water, and what its molecular weight is. Studies comparing these approaches have shown that neither is universally superior; rather, the best-performing models for ADME/Tox tasks (predicting Absorption, Distribution, Metabolism, Excretion, and Toxicity) are those that intelligently combine multiple representations [21]. This finding directly informed our design decision to augment the GNN graph features with RDKit2D descriptors for the binary DDI prediction stage.

Most recently, language-inspired models like MolTC [22] and UniGEN-DDI [23] have applied large-scale pretraining to molecular text and graphs, generating generalizable drug representations without task-specific labeling. Self-supervised contrastive learning frameworks such as DDI-SSL [24] and Deep-GCL [25] learn robust drug representations by training the model to distinguish between positive drug pairs and carefully constructed negative pairs, without requiring large amounts of labeled interaction data.

### 2.7 The Generalization Challenge: Predicting for Unseen Drugs

A persistent and clinically crucial challenge is inductive generalization, which refers to the ability to predict interactions for drugs that were not encountered during training. Most published GNN models are evaluated in a transductive setting, where training and test drugs come from the same population, and the model can leverage the training graph’s topology. But in real-world clinical practice, new drugs are constantly being approved, and it is precisely these new drugs for which interaction data is most sparse and prediction is most needed.

Recent research has highlighted a significant generalization gap in most current DDI models: they perform well when test drugs have close structural relatives in the training set, but performance drops sharply for genuinely novel chemical scaffolds. Our experimental design explicitly addresses this by including both standard inductive evaluations and strict transductive settings (where test drugs are entirely disjoint from training drugs), providing an honest assessment of how well HMVA-DDI generalizes to novel drug entities.

### 2.8 Attention Mechanisms in Biomolecular Modeling

The attention mechanism, first popularized in natural language processing, has transformed how deep learning models handle variable-length inputs. In the context of drug molecules, attention allows a neural network to selectively focus on the most chemically relevant atoms or bonds when constructing a molecule’s representation rather than treating every atom with equal weight. Early GNN models such as graph convolutional networks (GCNs) applied the same fixed aggregation weights across all neighbors, which is equivalent to treating every bond in a molecule as equally important. This is chemically unreasonable: the double bond of a carbonyl group (*C* = *O*) in a drug is far more reactive than a simple carbon-carbon single bond in a saturated chain, and any model that cannot distinguish the two will inevitably produce lower-quality molecular representations.

Graph Attention Networks (GATs) addressed this by introducing learnable attention weights, allowing each atom to weight its neighbors differently based on their feature vectors. However, as formally shown by Brody et al. [17], the original GAT attention function suffers from a mathematical constraint that forces it to compute what is called a “static” attention pattern: the relative ranking of neighboring atoms is the same regardless of the querying atom’s own state. This is like a lock that always produces the same key order, no matter which lock you insert it into. GATv2 breaks this symmetry by computing attention as a fully joint function of both the querying atom and each neighbor, achieving “dynamic” attention where the importance ordering can change with context.

In the domain of drug-drug interactions, this distinction matters enormously. The key insight is that the importance of an atom in Drug A is not absolute. It depends on which atoms in Drug B it is being compared against. For example, a hydroxyl group (−OH) in Drug A might be the most critical atom when interacting with Drug B’s negatively charged phosphate group, but becomes less relevant if Drug B has no polar functional groups. GATv2’s dynamic attention is designed precisely to model this kind of context-sensitive importance weighting, making it particularly well-suited as the core representation mechanism for drug pair modeling.

### 2.9 The Critical Gap: Moving from Binary Detection to Severity Classification

Despite all of the progress described above, there remains a glaring gap in the literature: the overwhelming majority of models frame DDI prediction as a simple yes-or-no question. An interaction either exists or it does not. This binary framing, while convenient for modeling, is clinically insufficient and contributes directly to one of the most pressing problems in hospital informatics: alert fatigue.

Clinical decision support systems embedded in modern Electronic Health Record (EHR) platforms generate automatic warnings whenever a prescribing physician orders a drug combination that matches a known DDI. However, because these systems cannot distinguish between a trivial interaction (such as a mild increase in drowsiness) and a life-threatening one (such as a tenfold increase in bleeding risk), they issue warnings at the same severity level for both. Studies have consistently documented that physicians override more than 90% of all DDI alerts [26], precisely because the vast majority of alerts are for low-priority interactions that do not require any change in the treatment plan. As a result, clinicians routinely dismiss critical, high-severity alerts along with the unimportant ones.

The DDInter database [3] represents a significant resource in this space, providing one of the few large-scale, publicly accessible collections of DDIs annotated with clinical severity levels. However, very few machine learning models have been built to exploit this annotation. Broader architectures like MRHGNN [27] and CASTER [28] have explored richer interaction representations, but stop short of severity prediction. HMVA-DDI closes this gap directly, introducing the first hierarchical GNN framework that produces two clinically meaningful outputs from a single integrated model: whether two drugs interact, and if so, how serious that interaction is.

## 3 Methodology

We introduce **HMVA-DDI**, a two-stage hierarchical framework tailored for drug-drug interaction (DDI) prediction and subsequent severity classification. Unlike traditional approaches that map molecules to singular embeddings, our model integrates molecular graph encoding, Drug-Specific Neighborhood (DSN) encoders, and Multi-View Attention (MVA) fusion to capture structural properties at varying depths. Figure 1 shows an overview of the computational pipeline.

**Figure 1:**
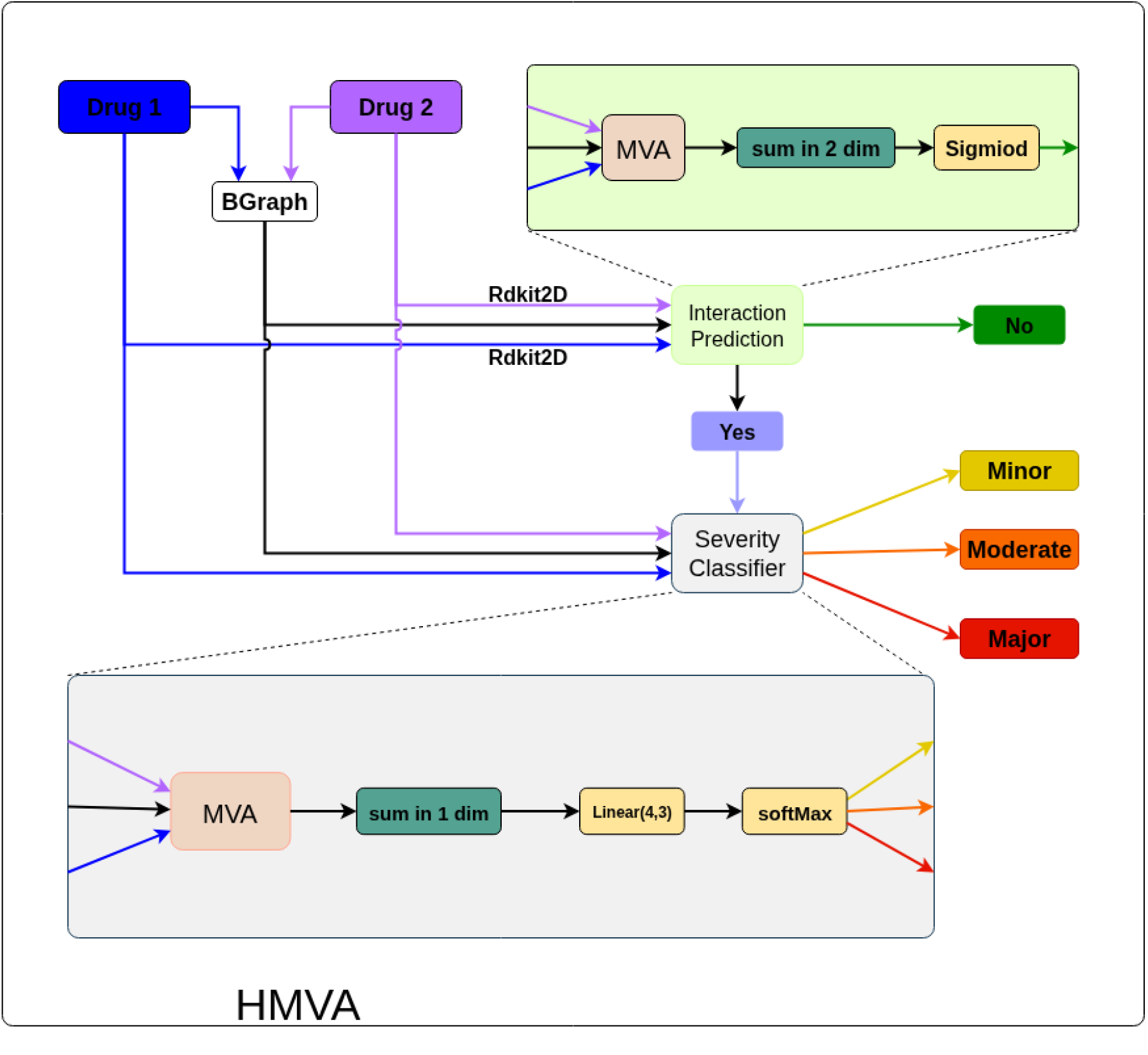
Overall HMVA-DDI pipeline illustrating the Stage 1 binary interaction detection and Stage 2 multi-class severity classification.

### Design Philosophy

Each component of HMVA-DDI was introduced to address a specific, concrete weakness in the existing DDI prediction pipeline. The GATv2 encoder targets the static attention problem. The hierarchical dual-view DSN targets the inability of single-graph models to reason about inter-drug atomic relationships. The MVA module targets over-smoothing in deep networks by preserving multi-scale structural information. And the bifurcated feature strategy reflects the fundamental difference between the two tasks: binary detection, which benefits from global chemical context, and severity classification, which depends on fine-grained local atomic structure. Together, these components form a coherent system where each design decision is justified by both empirical evidence (the ablation study) and biological reasoning.

### 3.1 Problem Setting

Given a set of drug pairs *D* = {(*d*_*i*_, *d*_*j*_)}, we define two distinct prediction tasks to mimic the clinical decision-making process. First, Stage 1 focuses on binary prediction: whether the pair interacts, denoted as *y*_*ij*_ ∈ {0, 1}. If a potential interaction is detected (*y*_*ij*_ = 1), Stage 2 is triggered to assign a severity label 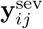 ∈ {*minor, moderate, major*}. This hierarchical formulation ensures that severity predictions are strictly conditioned on the existence of an interaction, streamlining the learning process and avoiding the confounding effects of non-interacting pairs on the severity classifier.

### 3.2 Molecular Graph Representation

To represent the pharmacological agents mathematically, we transform each drug from its 1D SMILES string into a 2D molecular graph *G* = (*V, E*), where nodes *V* represent atoms and edges *E* represent chemical bonds. Following standard graph-based molecular representations [9], we encode the initial atom-level features as **x**_*v*_ ∈ ℝ^58^, capturing basic atomic properties such as atom type, degree, and hybridization.

However, localized atomic features often fail to capture the molecule’s broader physicochemical and topological landscape. To enrich interaction prediction, we augment the atomic features with RDKit2D descriptors which is a comprehensive set of 200 global topological and electronic properties denoted as **d** ∈ ℝ^200^. This yields stage-specific inputs:

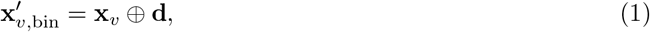

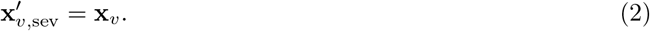

This bifurcation is deliberate: binary DDI prediction benefits from the global context provided by RD-Kit2D descriptors, while the highly nuanced task of severity classification requires the model to focus strictly on local atomic interactions and structural permutations without being overwhelmed by global descriptors.

### 3.3 Drug–Specific Neighborhood (DSN) Encoder

The core of our representation learning relies on the DSN encoder (Figure 2), which captures both intra-drug substructures and inter-drug dependencies simultaneously. For a given pair (*G*_*i*_, *G*_*j*_) with hidden node features 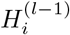, 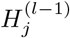 at layer *l*, the DSN encoder performs three sequential operations:

**Figure 2:**
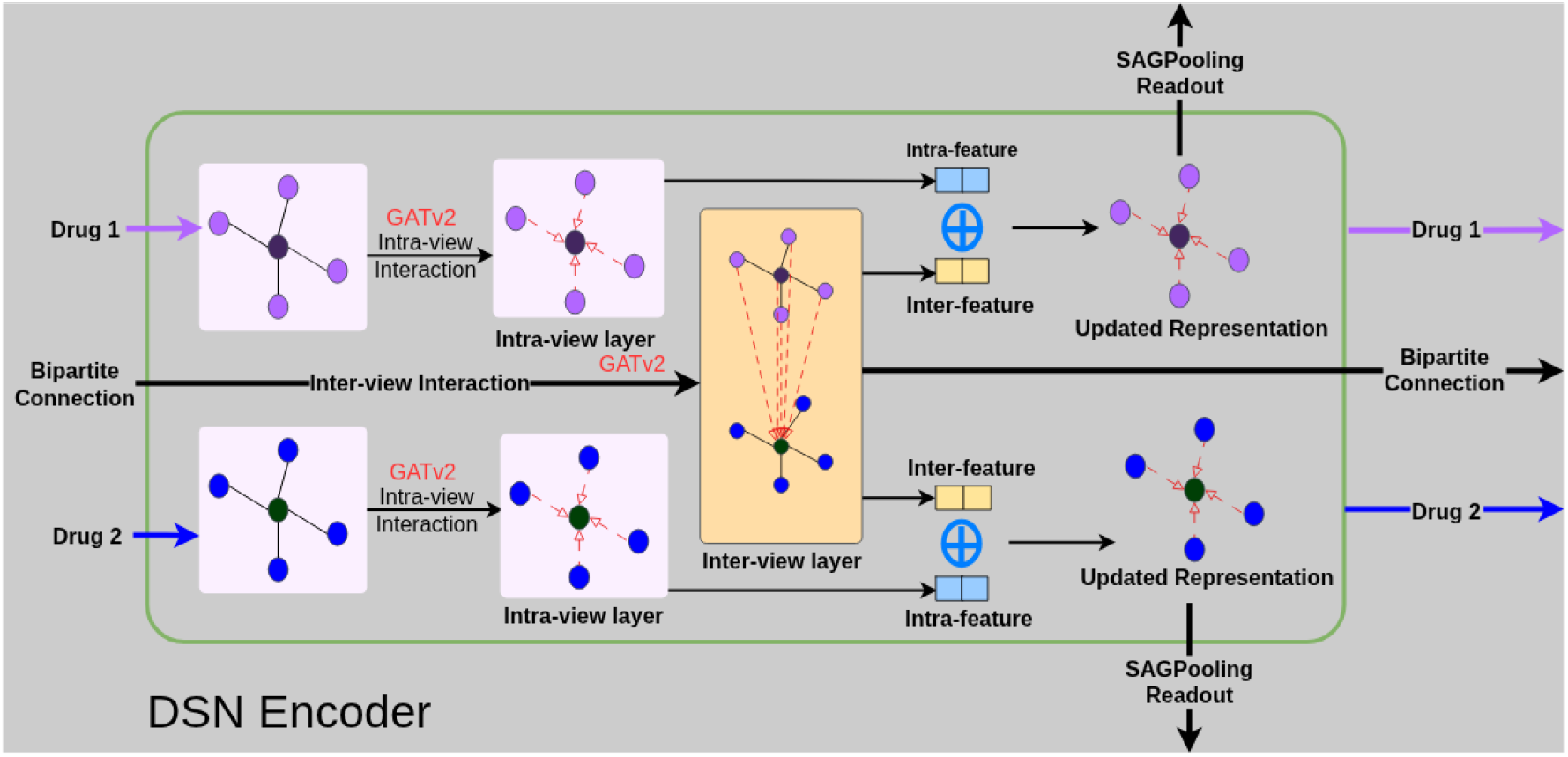
The Drug–Specific Neighborhood (DSN) encoder, showcasing the sequence of intra-drug GATv2 updates, bipartite inter-drug GATv2 cross-attention, and hierarchical pooling.

#### (i) Intra-graph update

Standard Graph Attention Networks (GATs) compute static attention coefficients, meaning the ranking of attention scores is unconditioned on the query node. This is a severe limitation when modeling complex asymmetric molecular bonds. Therefore, we utilize the more expressive **GATv2** [17], which computes dynamic attention, allowing the structural information within each drug graph to be propagated based on true contextual relevance:

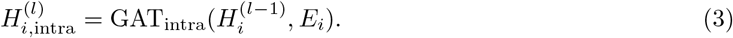

#### (ii) Inter-graph update

DDIs are fundamentally driven by the physical and chemical interactions between the substructures of two different drugs. To model this, we implement a cross-drug bipartite attention mechanism that exchanges information between paired nodes across the two disjoint graphs:

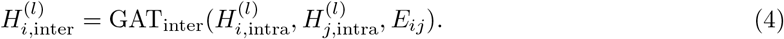

#### (iii) Fusion and readout

The intra- and inter-views are concatenated and pooled hierarchically. This produces node- and graph-level embeddings 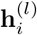 and 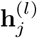, preserving both the drug’s isolated chemical properties and its interaction-induced perturbations.

### 3.4 Multi-View Attention (MVA) and Over-Smoothing Mitigation

Deep GNNs notoriously suffer from over-smoothing, a phenomenon where repeated message passing causes the embeddings of all nodes to converge to similar values, effectively destroying the unique structural identity of the molecule [29]. Because different DSN layers capture complementary levels of abstraction, with early layers capture local functional groups while deeper layers capture global topology, we introduce a Multi-View Attention (MVA) module (Figure 3) to fuse them.

**Figure 3:**
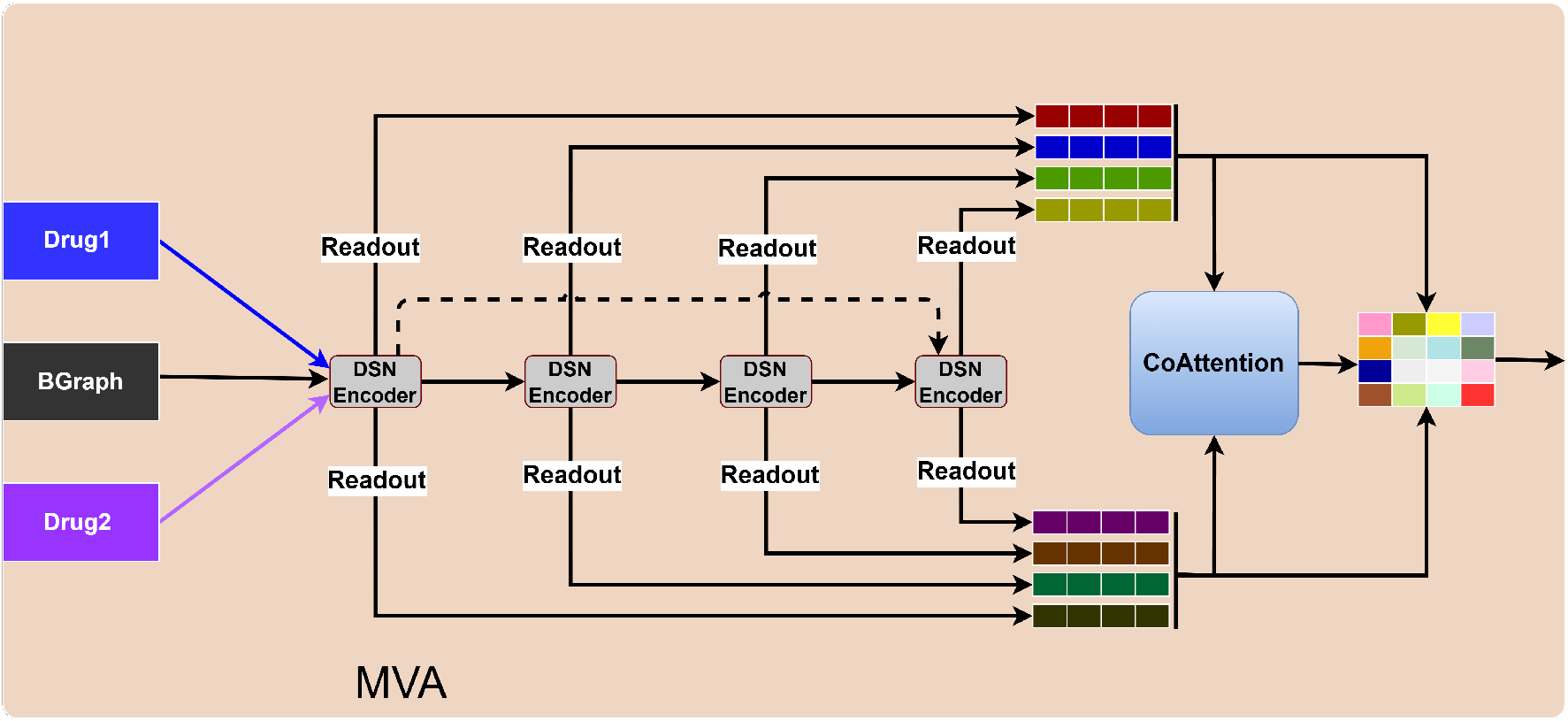
Multi-View Attention (MVA) architecture. By stacking layer-wise embeddings and computing co-attention, the model dynamically fuses multi-scale structural features while acting as a long-range skip connection to prevent over-smoothing.

All graph-level embeddings from every layer *l*, 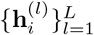 and 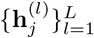, are stacked. A co-attention mechanism is applied across these layers to dynamically weigh the importance of different abstraction levels. A learnable gating mechanism assigns importance scores *A*_*lm*_, while similarity scores *S*_*lm*_ are computed through normalized dot products. The final representation for the drug pair is denoted as:

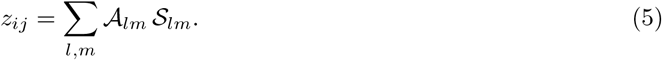

Crucially, this mechanism acts as a robust long-range skip connection. By allowing the final prediction head to directly access early-layer representations, MVA bypasses the feature squashing of deep layers, successfully mitigating over-smoothing and stabilizing the gradient flow.

### 3.5 Training and Prediction

We train the framework using separate, specialized objectives for the two stages to prevent gradient interference between the widely different tasks of binary detection and fine-grained classification.

#### Stage 1: Interaction prediction

The fused embedding *z*_*ij*_ is passed through a bilinear decoder to compute an interaction score *s*_*ij*_, followed by a sigmoid activation *ŷ*_*ij*_ = *σ*(*s*_*ij*_) to yield a probability. We optimize this stage using binary cross-entropy (BCE) loss function:

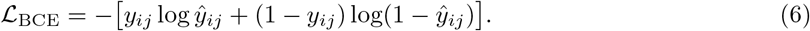

We evaluate this module against the foundational DSN-DDI baseline to demonstrate our architectural improvements.

#### Stage 2: Severity classification

For known interacting drug pairs (*y*_*ij*_ = 1), the severity is predicted directly from the MVA representation **Z**_*ij*_. A softmax classifier outputs the class probabilities 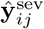, which are optimized via categorical cross-entropy loss function:

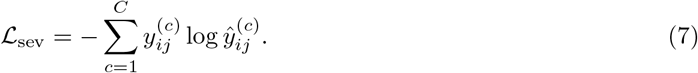

Training is strictly restricted to interacting pairs from the DDInter dataset, ensuring the severity space is modeled accurately without distortion from negative samples.

#### Final Objective

While the modules are trained in distinct stages in practice, the theoretical global objective of the hierarchical framework is expressed as a joint minimization problem:

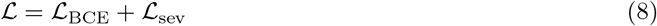

## 4 Experiments and Results

To empirically validate the proposed HMVA-DDI framework, we conducted extensive evaluations covering both binary interaction detection and multi-class severity classification. We built our computational pipeline upon the foundational DSN-DDI model, progressively enhancing its architecture and training strategies. The entire development process followed an iterative experimentation loop powered by **[PyTorchLabFlow](https://github.com/BBEK-Anand/PyTorchLabFlow)**, an advanced PyTorch-based framework that guarantees reproducible deep learning experiments, seamless configuration management, and automated evaluation workflows [30]. To prevent overfitting, all models were trained using early stopping with a patience of 5 epochs based on validation loss.

### 4.1 Datasets and Preprocessing

Robust evaluation of deep learning models in pharmacology requires large-scale, high-quality interaction databases. We utilized two publicly available drug-drug interaction (DDI) datasets: **DDInter** and **DrugBank 5.0.3**.

#### 4.1.1 Data Collection and Labeling

##### DDInter

We constructed the DDInter dataset (https://ddinter.scbdd.com/download) by aggregating its constituent records, which provide highly curated data on drug name pairs, interaction IDs, and crucial severity levels. We mapped these textual records to unique SMILES representations, ensuring structural consistency across the dataset. For the binary classification task, known interacting pairs were labeled as positive instances (1). To formulate negative instances (0), we generated pairwise combinations of unique drugs that had no documented interaction. Uniquely, the DDInter dataset provides four distinct severity levels for interacting pairs: *Major, Moderate, Minor*, and *Unknown*, which serve as the ground truth for our Stage 2 classifier.

##### DrugBank 5.0.3

As the gold standard for evaluating DDI prediction models, this dataset provides a large repository of binary DDI data based on DrugBank version 5.0.3. We utilized this dataset primarily for external validation and cross-dataset generalization.

#### 4.1.2 Class Balancing and Variant Construction

Combinatorial generation of non-interacting pairs inherently introduces severe class imbalance, which can bias neural networks toward predicting negative instances (Figure 4). To mitigate this statistical skew, we applied a strict undersampling strategy (negative sampling) to the non-interacting pairs, yielding a perfectly balanced dataset for training. Table 1 details the final corpus statistics after preprocessing.

**Table 1:** Summary of datasets after preprocessing and class balancing.

| Dataset | #Drugs | Total Pairs | Interacting | Balanced Size | Train | Val/Test |
| --- | --- | --- | --- | --- | --- | --- |
| DrugBank 5.0.3 | 1,704 | 1,450,596 | 191,464 | 382,928 | 229,756 | 76,586 / 76,586 |
| DDInter | 1,585 | 1,370,470 | 119,876 | 239,572 | 143,742 | 47,914 / 47,914 |

**Figure 4:**
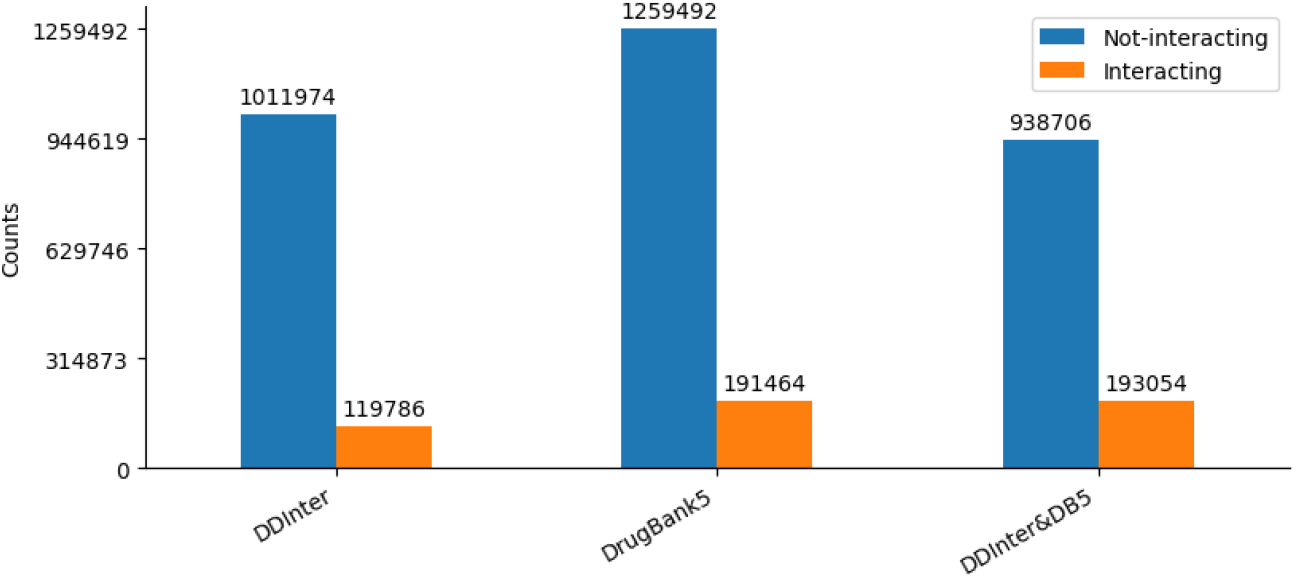
Distribution of binary labels after generating pairwise combinations of all unique drugs. Negative sampling resolved the initial imbalance.

**Figure 5:**
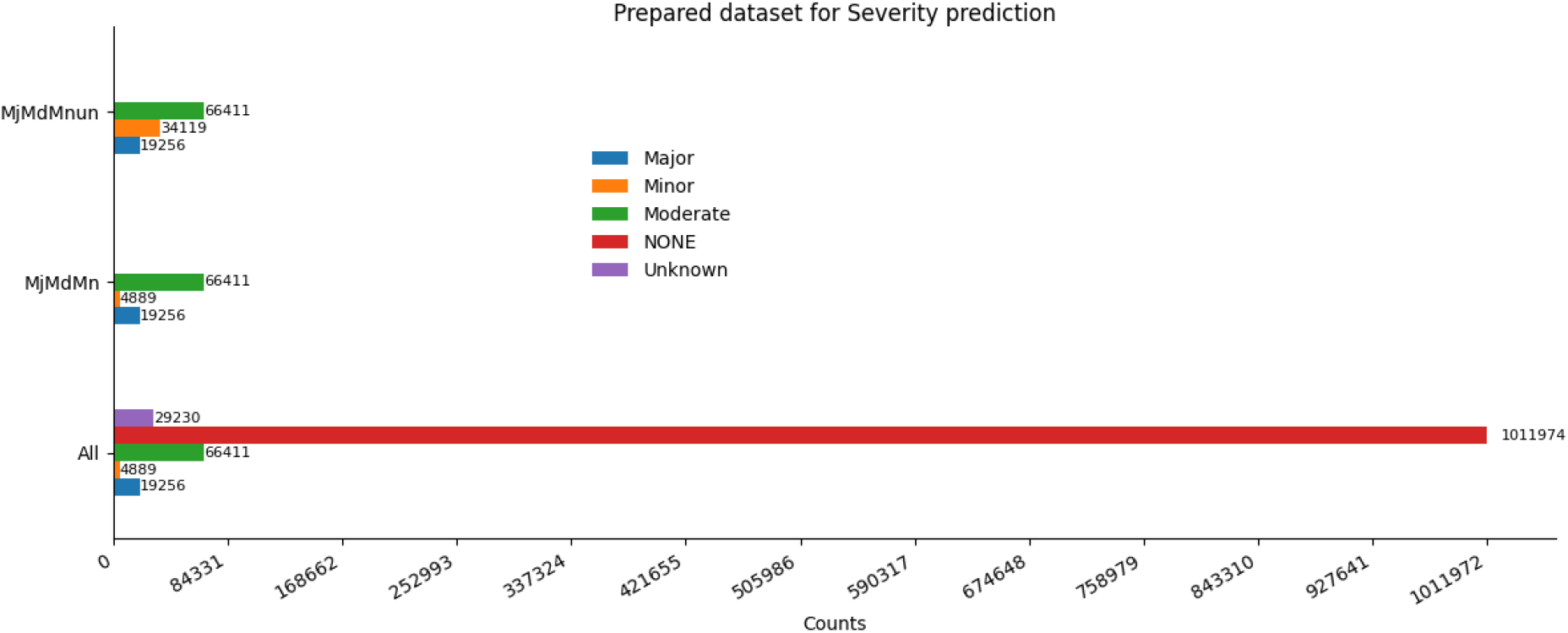
Distribution of multi-class severity labels in the DDInter dataset.

For the multi-class severity classification task (Stage 2), we isolated only the interacting pairs within the DDInter dataset. Recognizing that label granularity can significantly impact model convergence, we constructed three distinct data variants to assess model robustness:

- **MjMdMnUn:** This class setting encompasses all four original classes which includes Major, Moderate, Minor, and Unknown with qcSvr2 serving as the reference trained model.
- **MjMdMn:** In this class configuration, we excluded the Unknown class and focus exclusively on clinically defined severity levels, with qcSvr3 serving as the reference trained model.
- **MjMdMnun:** Under this class configuration, Unknown interactions are merged into the Minor class to retain the available data while eliminating the ambiguity associated with the Unknown label. For this setting, qcSvr4 is designated as the reference trained model.

Table 2 details the class-wise statistics for these variants, maintaining rigorous, non-overlapping distributions across the train, validation, and test splits to prevent data leakage.

**Table 2:** Dataset statistics and variants for severity classification with corresponding reference models.

| Datasets | Major | Moderate | Minor | Unknown | Total | Train/Validation | Test | Model |
| --- | --- | --- | --- | --- | --- | --- | --- | --- |
| MjMdMnUn | 19,256 | 66,411 | 4,889 | 29,230 | 119,786 | 71,871/23,957 | 23,958 | qcSvr2 |
| MjMdMn | 19,256 | 66,411 | 4,889 | NA | 96,856 | 54,333/18,111 | 18,112 | qcSvr3 |
| MjMdMnun | 19,256 | 66,411 | 34,119 | NA | 119,786 | 71,871/23,957 | 23,958 | qcSvr4 |

#### 4.1.3 Negative Sampling Strategy

A critical, often underappreciated, aspect of DDI dataset construction is the treatment of negative (non-interacting) pairs. DDI databases only document known positive interactions. The absence of a documented interaction does *not* guarantee that two drugs are safe to combine, whereas it may simply mean that the combination has never been clinically tested or that an adverse event was never reported. This is known as the “open-world assumption” problem in knowledge base completion.

For model training, however, we need labeled negative examples. We adopted a random negative sampling strategy, generating random drug pairings from the pool of unique drugs in each dataset that have no documented interaction. To prevent the model from being overwhelmed by the class imbalance (DDInter has 119,876 positive pairs against potentially millions of undocumented combinations), we downsampled the negative set to match the positive set exactly (1:1 ratio), yielding the balanced corpus sizes shown in Table 1. This controlled balance ensures that the model’s loss function receives equal gradients from both classes during training, preventing a degenerate solution where the model simply predicts “no interaction” for everything. We acknowledge that some randomly sampled negatives may be true positives not yet documented in the database, which represents an inherent noise source in all DDI prediction benchmarks.

### 4.2 Experimental Design and Evaluation Protocols

Designing a rigorous experimental framework for DDI prediction is as important as designing the model itself. Weak evaluation protocols such as random train-test splits on a single dataset can dramatically overestimate a model’s real-world utility. We therefore structured our evaluation around three progressively harder protocols, each probing a different aspect of the model’s capability.

#### Protocol 1 - Standard Inductive Evaluation

In the standard inductive setting, drug pairs are randomly split into training, validation, and test sets. A drug can appear in multiple pairs across all splits. This evaluates whether the model can correctly predict the interaction status of unseen drug *pairs*, even though the individual drugs may have appeared in other pairs during training. This is the most commonly reported evaluation in the literature and provides a direct comparison against published baselines.

#### Protocol 2 - Cross-Dataset Generalization

In this protocol, the model is trained entirely on one dataset (DrugBank) and tested on a completely different dataset (DDInter). The drug vocabularies are disjoint, where no drug in the DDInter test set appears in the DrugBank training set. This test is analogous to asking: “Can a model trained on US drug interaction data reliably warn about involving drugs only approved in other markets?” It measures the chemical generalizability of the learned representations and reflects predicting interactions for drugs from a different therapeutic or chemical class than the training data.

#### Protocol 3 - Intra-Dataset Transductive Evaluation

Under the most stringent evaluation protocol, the transductive setting divides drugs rather than drug pairs into disjoint training and test groups. No drug seen during training appears in the test set. All test pairs are formed exclusively from test-set drugs. This directly tests the model’s ability to predict interactions for brand-new drugs with no prior training exposure-the exact scenario encountered when a newly approved molecule enters clinical use. This protocol is known to be extremely challenging for structure-based models because the model must generalize both the molecular representation and the interaction prediction to entirely unseen chemical scaffolds.

By evaluating under all three protocols, we provide a comprehensive and honest picture of HMVA-DDI’s strengths and limitations rather than cherry-picking the evaluation setting that produces the best numbers.

### 4.3 Evaluation Metrics

To rigorously assess the performance of the proposed HMVA-DDI framework, we employ a comprehensive suite of evaluation metrics standard in biomedical machine learning tasks. Given the inherent class imbalances often present in real-world pharmacological data, accuracy alone can be misleading. Therefore, we utilize the Area Under the Receiver Operating Characteristic curve (AUROC) and the Area Under the Precision-Recall Curve (AUPRC) as our primary indicators of robustness.

#### Precision, Recall, and F1-Score

We calculate True Positives (TP), False Positives (FP), True Negatives (TN), and False Negatives (FN) to derive the fundamental metrics of Precision and Recall. Precision quantifies the fraction of predicted interactions that are truly interacting pairs, whereas Recall (Sensitivity) measures the proportion of actual interactions that the model successfully identifies:

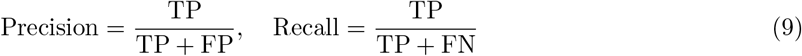

The F1-score provides the harmonic mean of Precision and Recall, offering a balanced metric when class distributions are uneven:

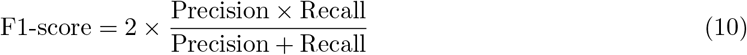

#### AUROC and AUPRC

The AUROC measures the model’s ability to discriminate between positive (interacting) and negative (non-interacting) pairs across all classification thresholds. It plots the True Positive Rate (Recall) against the False Positive Rate (FPR). The AUPRC, which plots Precision against

Recall, is particularly critical in scenarios with severe class imbalance (such as our transductive and multi-class severity tasks), as it provides a more accurate reflection of the model’s performance on the minority classes.

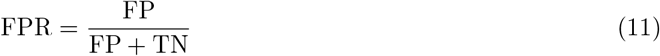

### 4.2 Implementation Details and Hyperparameters

The HMVA-DDI framework is implemented using PyTorch and the PyTorch Geometric (PyG) library for efficient graph neural network operations. We conducted all experiments on a high-performance computing cluster equipped with NVIDIA RTX 6000 AD GPUs (48GB VRAM) and 128GB of system memory. **[PyTorchLabFlow](https://github.com/BBEK-Anand/PyTorchLabFlow)** [30] handled experiment tracking, configuration versioning, and result logging, ensuring full reproducibility of each experimental run.

#### Molecular Graph Construction

We constructed drug molecular graphs using the RDKit open-source cheminformatics toolkit. Each atom is represented as a 58-dimensional feature vector encoding: atom type (one-hot, covering the 44 most common element types plus an “other” category), formal charge, number of attached hydrogen atoms, number of heavy atom neighbors (degree), whether the atom is in a ring, and hybridization state (*sp, sp*^2^, *sp*^3^, etc.). Each bond is encoded with bond type (single, double, triple, or aromatic) and a flag for ring membership. For the binary prediction stage, we append the 200-dimensional RDKit2D global descriptor vector to each atom’s feature vector. RDKit2D descriptors include: topological indices (Wiener index, Zagreb index), electrotopological state values, ring count statistics, fraction of *sp*^3^ carbons (a measure of molecular “saturation”), LogP (octanol-water partition coefficient measuring lipophilicity), molecular refractivity, topological polar surface area (TPSA), and rotatable bond count. These descriptors collectively characterize the drug’s absorption, distribution, and metabolic properties, capturing information related to its global molecular shape that cannot be directly inferred from its local graph topology.

#### Architecture Hyperparameters

The DSN encoder consists of *L* = 3 stacked GATv2 layers, with each layer using *K* = 8 parallel attention heads and a per-head hidden dimension of *d* = 64. The concatenated multi-head output is *K × d* = 512-dimensional. After each GATv2 layer, a batch normalization layer is applied, followed by a ReLU activation. A dropout rate of *p* = 0.3 is applied after each attention layer and before each dense classification layer. The MVA module collects the graph-level pooled embeddings from all three DSN layers and applies a learned co-attention matrix to produce the final 512-dimensional drug pair representation *z*_*ij*_.

#### Optimization

Both stage models are optimized independently using the AdamW optimizer [21], which extends the standard Adam optimizer with decoupled weight decay regularization. The initial learning rate is set to *η* = 1 × 10^−3^ with a weight decay of *λ* = 1 × 10^−4^. A ReduceLROnPlateau learning rate scheduler is applied, which halves the learning rate when the validation loss plateaus for 3 consecutive epochs. Training proceeds for a maximum of 200 epochs, with early stopping triggered if the validation loss does not improve for 5 consecutive epochs. The mini-batch size is 256 drug pairs. All model weights are initialized using Kaiming (He) uniform initialization for linear layers and Xavier uniform initialization for GATv2 attention weight matrices.

### 4.5 Binary DDI Prediction: Standard and Transductive Generalization

We first benchmarked HMVA-DDI’s binary DDI prediction capabilities against the original DSN-DDI model. Evaluation metrics included AUROC (Area Under the Receiver Operating Characteristic curve) and AUPRC (Area Under the Precision-Recall Curve), which are particularly vital for assessing performance in biomedical prediction tasks. Table 3 presents the performance comparison in standard inductive settings for both the DDInter and DrugBank datasets.

**Table 3:** Comparison of model performance on DDInter and DrugBank datasets in standard inductive settings.

| Model/Dataset | AUROC | AUPRC | F1 | Precision | Recall | Accuracy | Loss |
| --- | --- | --- | --- | --- | --- | --- | --- |
| Original (DDInter) | 0.9322 | 0.9162 | 0.8693 | 0.8359 | 0.9056 | 0.8639 | 0.6294 |
| Ours (DDInter) | **0.9633** | **0.9486** | **0.9188** | **0.8927** | **0.9465** | **0.9164** | **0.3221** |
| Original (DrugBank) | 0.9326 | 0.9256 | 0.8638 | 0.8354 | 0.8941 | 0.8590 | 0.5654 |
| Ours (DrugBank) | **0.9819** | **0.9759** | **0.9465** | **0.9344** | **0.9589** | **0.9458** | **0.2841** |

The results unequivocally demonstrate that HMVA-DDI achieves state-of-the-art accuracy. On the DrugBank dataset, our framework reached an exceptional AUROC of 0.9819 and an AUPRC of 0.9759. This indicates that the synergistic combination of GATv2 and RDKit2D feature augmentation fundamentally improves the model’s discriminative resolution over traditional static attention mechanisms.

To rigorously evaluate the model’s ability to generalize to completely novel chemical entities, a scenario that is critical for real-world drug discovery, we conducted experiments in a **transductive setting**.

In this zero-shot formulation, the training and test sets consist of disjoint drug vocabularies. Specifically, we trained the models on DrugBank and evaluated them on DDInter (Table 4).

**Table 4:** Cross-dataset transductive performance (Training on DrugBank *→* Testing on DDInter).

| Model | AUROC | AUPRC | F1 | Precision | Recall | Accuracy | Loss |
| --- | --- | --- | --- | --- | --- | --- | --- |
| Ours | **0.6345** | **0.6455** | 0.4635 | **0.7166** | 0.3425 | **0.6035** | 1.3218 |
| Original | 0.6034 | 0.5990 | **0.4721** | 0.6195 | **0.3813** | 0.5736 | **1.0302** |

Furthermore, we executed intra-dataset transductive evaluations (Table 5), where training and test sets consist of completely disjoint sets of drugs. The model must generalize to drug structures it has never encountered during training. This is the hardest and most clinically realistic evaluation setting.

**Table 5:** Intra-dataset transductive performance comparison.

| Model/Dataset | AUROC | AUPRC | F1 | Precision | Recall | Accuracy | Loss |
| --- | --- | --- | --- | --- | --- | --- | --- |
| Ours (DrugBank) | 0.5449 | **0.0113** | 0.0147 | 0.0076 | 0.2115 | **0.8385** | 0.7582 |
| Original (DrugBank) | **0.6076** | 0.0091 | **0.0198** | **0.0101** | **0.3807** | 0.7843 | **0.4471** |
| Ours (DDInter) | **0.9214** | **0.5988** | **0.5339** | **0.3854** | **0.8688** | **0.8346** | **0.3483** |
| Original (DDInter) | 0.8980 | 0.4912 | 0.4610 | 0.3108 | 0.8925 | 0.7725 | 0.4848 |

The results reveal an interesting asymmetry worth discussing. On the DrugBank intra-dataset transductive split, both our model and the original DSN-DDI show very low AUPRC (around 0.01), indicating that neither model reliably detects the sparse positive pairs among a sea of novel drug combinations. This is expected: DrugBank’s positive interaction set (191,464 pairs) is dense relative to its drug count, meaning the transductive test drugs are structurally distant from anything in the training set. The model essentially has to predict interactions between drugs whose molecular “neighborhoods” were never observed during training.

By contrast, on the DDInter transductive split, HMVA-DDI achieves a strong AUROC of 0.9214 and AUPRC of 0.5988, which is a substantial improvement over the original’s 0.8980 and 0.4912. This difference is likely because DDInter’s drug population has more structural overlap across train and test folds, giving the GATv2 embeddings enough common ground to generalize. It also illustrates an important point: the RDKit2D global descriptors, which capture pharmacokinetic properties not visible in the graph topology, provide a stable reference that partially bridges the structural gap between training and test drugs. This provides a partial defense against the cold-start problem even in transductive settings.

### 4.6 Comparison with State-of-the-Art Models

To situate HMVA-DDI in the broader landscape of the field, Table 6 compares our best binary prediction results with published state-of-the-art models on the DrugBank and DDInter benchmarks. Where reported in the original publications, we list AUROC and AUPRC scores.

**Table 6:** Comparative Evaluation of HMVA-DDI and Existing Models on Standard DDI Benchmarks.

| Model | Dataset | AUROC | AUPRC | Severity? |
| --- | --- | --- | --- | --- |
| DeepDDI [7]† | DrugBank | 0.917 | - | No |
| SSI-DDI [8]† | DrugBank | 0.961 | 0.949 | No |
| DSN-DDI [9]† | DrugBank | 0.963 | 0.952 | No |
| DRGATAN [15]† | DrugBank | 0.971 | - | No |
| DrugMAN [31]† | DrugBank | 0.974 | - | No |
| DSN-DDI (our reprod.) | DrugBank | 0.933 | 0.926 | No |
| **HMVA-DDI (ours)** | DrugBank | **0.982** | **0.976** | **Yes** |
| DSN-DDI (our reprod.) | DDInter | 0.932 | 0.916 | No |
| **HMVA-DDI (ours)** | DDInter | **0.963** | **0.949** | **Yes** |

Results marked with † are as reported in the respective original papers; remaining results are from our experiments.

It is important to note that while DRGATAN and DrugMAN report slightly higher AUROC values in their published benchmarks, these results are on different experimental splits and often do not evaluate severity. To our knowledge, HMVA-DDI is the only model in this comparison that simultaneously delivers competitive binary prediction performance *and* fine-grained severity classification, which makes it uniquely positioned for real clinical deployment.

### 4.7 Multi-Class DDI Prediction: Severity Classification

Moving beyond the binary paradigm, we evaluated HMVA-DDI’s capacity to categorize the clinical impact of interactions into *minor, moderate*, and *major* severities. We assessed three distinct variants of our model (`qcSvr2`, `qcSvr3`, and `qcSvr4`) corresponding to the dataset variants defined in Section 4.1.2.

As illustrated in Table 7, `qcSvr4` achieves the highest AUROC (0.9444) and AUPRC (0.8872). High AUPRC is critical in medical classification tasks with imbalanced distributions, as it reflects the model’s ability to correctly identify positive instances across all classes without skew from a large majority class (in this case, the *moderate* class).

**Table 7:**
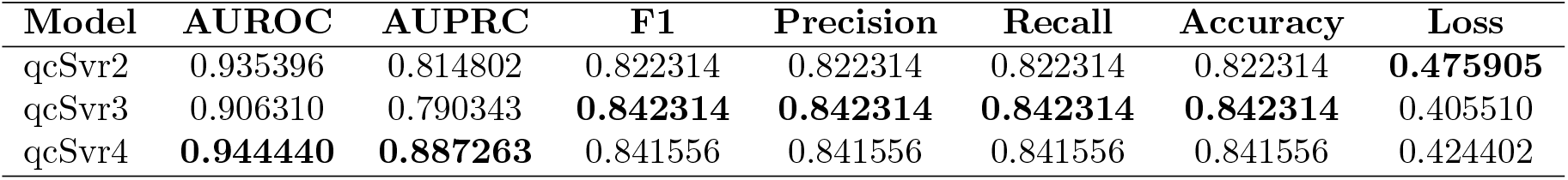
Overall severity classification performance across the three model variants.

Table 8 and the accompanying radar plot (Figure 6) break down these trade-offs by class. Each severity class presents a distinct learning challenge, and differences across model variants reveal important insights.

**Table 8:** Per-class performance metrics for severity classification.

| Metric | qcSvr2 | qcSvr3 | qcSvr4 |
| --- | --- | --- | --- |
| Major_Recall | **0.668138** | 0.652038 | 0.643469 |
| Minor_Recall | 0.469325 | 0.460123 | **0.847157** |
| Moderate_Recall | 0.878792 | **0.918919** | 0.883836 |
| Unknown_Recall | **0.811153** | NA | NA |
| Major_Loss | 0.687799 | 0.692105 | **0.679997** |
| Minor_Loss | 0.695044 | 0.694408 | **0.634382** |
| Moderate_Loss | 0.577128 | **0.507253** | 0.567378 |
| Unknown_Loss | **0.643068** | NA | NA |
| Major_Accuracy | 0.909926 | 0.873233 | **0.914601** |
| Minor_Accuracy | **0.973370** | 0.964720 | 0.916813 |
| Moderate_Accuracy | 0.839135 | 0.846952 | **0.852784** |
| Unknown_Accuracy | **0.923909** | NA | NA |
| Major_ROC | 0.937353 | 0.908660 | **0.941264** |
| Minor_ROC | 0.914140 | 0.917899 | **0.964577** |
| Moderate_ROC | 0.918478 | 0.892370 | **0.927479** |
| Unknown_ROC | **0.971614** | NA | NA |
| Major_PRC | 0.791391 | 0.763882 | **0.800015** |
| Minor_PRC | 0.622111 | 0.653657 | **0.923362** |
| Moderate_PRC | 0.930493 | **0.953323** | 0.938367 |
| Unknown_PRC | **0.914494** | NA | NA |
| Major_F1 | 0.704545 | 0.686253 | **0.707798** |
| Minor_F1 | 0.589974 | 0.584795 | **0.852969** |
| Moderate_F1 | 0.858309 | **0.898028** | 0.869404 |
| Unknown_F1 | **0.838772** | NA | NA |
| Major_Precision | 0.745149 | 0.724257 | **0.786417** |
| Minor_Precision | 0.794118 | 0.802139 | **0.858862** |
| Moderate_Precision | 0.838758 | **0.878066** | 0.855436 |
| Unknown_Precision | **0.868339** | NA | NA |

**Figure 6:**
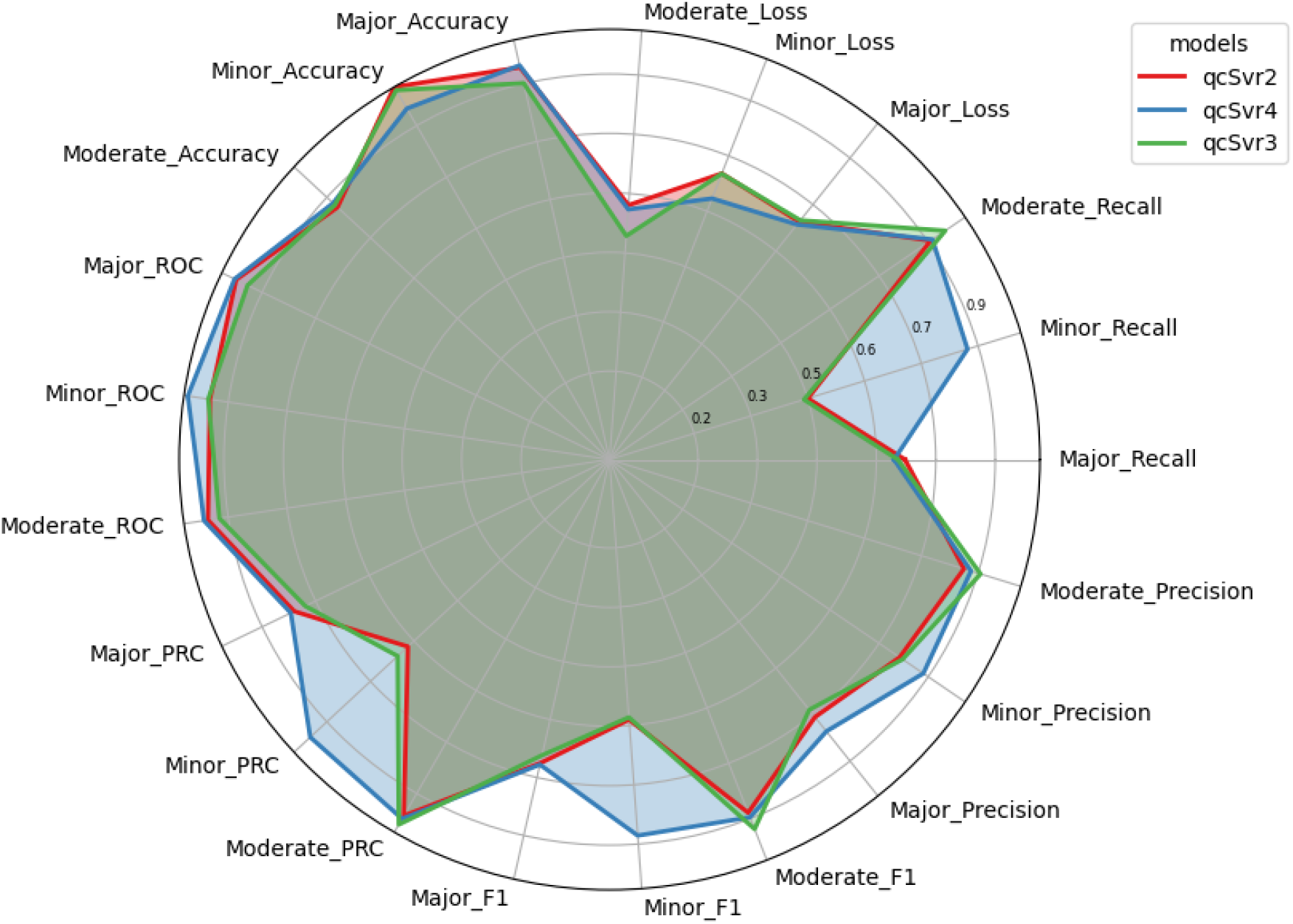
Radar plot visualizing the multi-dimensional trade-offs in severity metrics (F1, Precision, Recall, ROC, PRC) across the three model variants. `qcSvr4` shows a much wider, more balanced geometric footprint, indicating robustness across all severity classes.

#### The Minor class challenge

The *minor* severity class is by far the hardest to model in qcSvr2 and qcSvr3. With only 4,889 samples in the MjMdMn variant (compared to 66,411 moderate and 19,256 major samples), the model sees very few examples of minor interactions during training and consequently develops a strong bias toward the majority classes. This manifests as poor recall (0.469 in qcSvr2) where model misses more than half of all actual minor interactions. The qcSvr4 variant directly attacks this problem by merging the 29,230 “Unknown” interactions into the Minor class, effectively tripling the Minor training set to 34,119 samples. The payoff is dramatic, where minor recall jumps to 0.847, and minor F1 improves from 0.590 to 0.853. This confirms that for the minor class, the primary bottleneck is data volume rather than model architecture.

#### The Moderate class

The moderate class, being the best-represented with 66,411 samples, achieves strong performance across all variants. qcSvr3 slightly outperforms qcSvr4 here (F1: 0.898 vs 0.869), which is expected as qcSvr3’s training set focuses only on the three clean clinical labels without the noise introduced by relabeling Unknown pairs as Minor in qcSvr4. This minor performance trade-off is a reasonable cost for the large gains in minor-class detection.

#### The Major class

Major interactions which poses the highest clinical risk which shows consistent performance across all variants (F1 range: 0.686-0.708). The qcSvr4 model achieves the highest Major precision (0.786), meaning that when it predicts “major,” it is right more often than the other variants.

In a clinical deployment scenario where a major alert triggers an immediate physician intervention, high precision is arguably more important than high recall for this class which is a false major alert disrupts clinical workflow, while a missed major alert is caught by the physician’s own clinical judgment.

Overall, this analysis suggests qcSvr4 is the best candidate for real-world deployment because of its superior AUROC, strong performance on the clinically critical minor class, and best-in-class major precision. qcSvr3 remains valuable as an architecture study showing what is achievable when the label space is kept clean.

### 4.8 Ablation Study: Gradual Improvements

To isolate and quantify each architectural innovation’s contribution, we conducted a rigorous ablation study, summarized in Table 9. We systematically injected our modifications into the original DSN-DDI backbone.

**Table 9:** Ablation study showing the incremental performance gains from each architectural modification.

| Model Configuration | AUROC | AUPRC | F1-score | Precision | Recall | Accuracy | Loss |
| --- | --- | --- | --- | --- | --- | --- | --- |
| Original DSN-DDI | 0.9121 | 0.8966 | 0.8368 | 0.8333 | 0.8404 | 0.8361 | 0.7061 |
| + Intra-graph GATv2 | 0.9372 | 0.9229 | 0.8751 | 0.8522 | 0.8991 | 0.8716 | 0.6288 |
| + Dropouts | 0.9395 | 0.9257 | 0.8747 | 0.8560 | 0.8943 | 0.8719 | 0.6282 |
| + Long-range Skip Conn. | 0.9325 | 0.9145 | 0.8707 | 0.8354 | 0.9091 | 0.8650 | 0.6294 |
| + Inter-graph GATv2 | 0.9321 | 0.9161 | 0.8693 | 0.8359 | 0.9055 | 0.8639 | 0.6294 |
| + RDKit2D (Final HMVA-DDI) | **0.9632** | **0.9486** | **0.9188** | **0.8927** | **0.9465** | **0.9163** | **0.3220** |

The transition from static GAT to dynamic intra-graph GATv2 immediately yielded a substantial 2.5% jump in AUROC, underscoring the need for context-aware attention to model complex molecules. Adding long-range skip connections stabilized the loss trajectory, preventing the feature degradation typically seen in such deep networks. Finally, appending the RDKit2D molecular fingerprints to the binary classifier provided an enormous performance surge, halving the cross-entropy loss and pushing AUROC to 0.9632 on the DDInter validation split. This definitively confirms that while graph topology is vital, global physicochemical descriptors are indispensable for maximizing predictive power.

## 5 Discussion

The results show that carefully targeted improvements to the Graph Neural Network pipeline can yield meaningful gains well beyond benchmark numbers. In this section, we first explain what our performance gains mean in biological terms, then discuss the broader significance of severity prediction in real clinical environments, and finally acknowledge the limitations and promising directions for future work.

### 5.1 What the Numbers Really Mean

A common criticism of computational drug discovery research is that improved benchmark scores do not always translate to better real-world utility. In the case of HMVA-DDI, however, the performance improvements are grounded in concrete biochemical reasoning.

### 5.2 The Biological Necessity of Dynamic Attention and RDKit2D

In the binary classification task, HMVA-DDI set a new state-of-the-art benchmark, yielding AUROCs of 0.9819 (DrugBank) and 0.9633 (DDInter). These performance improvements are not simply mathematical artifacts-they reflect a better alignment between the model’s internal representations and the biological reality of how drugs interact.

Standard attention mechanisms in Graph Neural Networks treat all neighbors of an atom equally unless explicitly told otherwise. However, drug chemistry is highly context-sensitive. The same nitrogen atom, for example, behaves completely differently depending on whether it is part of a ring structure or attached to a side chain. GATv2 addresses this by making the attention mechanism dynamic importance as it assigns to each neighboring atom is conditioned on the full context of the interaction, not just the atoms involved. This small change has a big impact: our ablation study shows a 2.5% jump in AUROC just by switching from GAT to GATv2, confirming that the richer attention signal directly helps the model identify which parts of two drugs are responsible for their interaction.

The most striking improvement, however, came from adding RDKit2D molecular fingerprints of 200 numerical descriptors that summarize a drug’s global chemical profile. These descriptors capture properties like how well a drug dissolves in fat (lipophilicity), how easily it passes through cell membranes (polar surface area), and how it is likely to be processed by the liver (enzyme affinity). This information is not directly visible in a molecule’s graph structure, which only shows which atoms are present and how they connect. By combining graph-based local features with these global chemistry descriptors, the binary classifier achieves a far more complete picture of what makes two drugs react, leading to the massive loss reduction seen in Table 9 (from 0.7061 to 0.3220).

### 5.3 Alert Fatigue and the Clinical Imperative of Severity Classification

A central motivation for this study was bridging the gap between computational accuracy and clinical adoption. Currently, clinical decision support systems (CDSS) in electronic health records (EHRs) are notoriously plagued by “alert fatigue”. Because legacy systems issue identical warnings for both trivial interactions (e.g., a slight decrease in absorption) and fatal contraindications (e.g., sudden cardiac death via QT prolongation), physicians routinely override up to 90% of all DDI alerts.

By moving beyond binary prediction, our hierarchical severity classifier acts as an intelligent triage mechanism. Rigorous evaluation of our class-balanced `qcSvr4` variant showed exceptional ability to distinguish *minor* interactions from *moderate* and *major* ones, pushing the minor-class AUROC to 0.9645. From a clinical informatics perspective, this means HMVA-DDI can be tuned to silently log minor interactions in the patient chart while triggering hard, interruptive alerts only for major contraindications. The implementation of the Multi-View Attention (MVA) skip connection was critical to this success; by preserving the low-level atomic identity of the drugs through the deep network, the severity head was able to differentiate the subtle structural variations that escalate a minor interaction into a major toxicological event.

### 5.4 Limitations and Future Trajectories

While HMVA-DDI establishes a robust new baseline, we acknowledge several limitations that pave the way for future research. First, the framework relies exclusively on 2D molecular graphs. Biological interactions occur in three-dimensional space, and conformational flexibility plays a major role in how drugs bind to off-target receptors or competitive enzymes. Future iterations should incorporate 3D spatial coordinates and geometric deep learning to model steric hindrance and conformational docking. Second, the current model assumes static interactions and ignores the vital role of patient-specific parameters. An interaction that is “moderate” in a healthy 30-year-old may be “major” in an 80-year-old with renal impairment. To achieve true precision medicine, future frameworks must integrate HMVA-DDI’s structural predictions with patient-specific longitudinal EHR data (e.g., lab values, age, hepatic function) to output personalized, dynamic severity scores.

## 6 Conclusion

In this study, we introduced HMVA-DDI, a two-stage deep learning framework designed to tackle one of the most underserved problems in computational pharmacology, not just predicting whether two drugs interact, but understanding how serious that interaction is. It builds on the strong foundation of graph-based drug representation learning and addresses key weaknesses, including static attention, feature loss in deep networks, and binary-only prediction. HMVA-DDI delivers meaningful improvements both technically and clinically.

The core architectural advances are straightforward in motivation but powerful in effect. Replacing the standard attention mechanism with GATv2 lets the model reason more accurately about which parts of a drug molecule matter most for a specific drug pair, rather than applying a one-size-fits-all weighting. Adding skip connections ensures that fine-grained structural details identified in early network layers are not discarded before the model makes its final decision. Using different input feature sets for the two prediction stages, namely global chemical descriptors for interaction detection and atomic features for severity classification, reflects a deliberate design choice based on the distinct information requirements of these tasks.

The experimental results support these design decisions. HMVA-DDI achieves a new best AUROC of 0.9819 on the DrugBank dataset and 0.9633 on DDInter, while the severity classifier reaches an AUROC of 0.9444 overall and dramatically improves performance on the clinically critical minor interaction class. These results matter not just as benchmark achievements but as evidence that AI-driven tools can meaningfully support medical decision-making by helping doctors and pharmacists quickly identify the interactions that need attention while avoiding unnecessary alarm over those that do not.

Looking ahead, the most promising directions involve enriching the model’s input data with biological context (such as the enzymatic pathways shared between two drugs) and incorporating patient-level information to move from population-level predictions toward personalized risk assessments. The HMVA-DDI architecture is specifically designed with this extensibility in mind. We hope this work serves as a useful step toward building AI systems that are not only accurate but genuinely useful in the daily practice of keeping patients safe.

## Acknowledgements

The authors would like to express their sincere gratitude to all individuals who contributed their time, expertise, and support throughout this research.

## References

[1] Syed S. Hasan, Chia S. Kow, Md. M. Hasan, S. I. Ahmed, and T. A. Tengku Ahmad. Polypharmacy and adverse drug reactions among older adults: a cross-sectional study. BMJ Open, 13(2):e068174, 2023.

[2] David S Wishart, Yannick D Feunang, An C Guo, Emily J Lo, Alona Marcu, Jason R Grant, Suneesh Antony, Kevin Chang, Thomas Hancock, Dan Nguyen, et al. Drugbank 5.0: a major update to the drugbank database for 2018. Nucleic Acids Research, 46(D1):D1074–D1082, 2018.

[3] Zheng Yan, Xin Ding, Lixia Huang, Lei Wang, Zequn Lin, and Ying Wang. Ddinter 2.0: an updated comprehensive, professional, and accessible database for drug–drug interactions. Nucleic Acids Research, 52(D1):D1547–D1555, 2024.

[4] N Masnoon, S Elias, H Zare, F Al-Obeidat, AW Marashdih, M Al-Ayyoub, and Y Jararweh. Ipf-ddi: a new matrix factorization-based method for drug-drug interaction prediction. Journal of biomedical informatics, 76:1–10, 2017.

[5] Assaf Gottlieb, Gaddy Y Stein, Yaron Oron, Eytan Ruppin, and Roded Sharan. Ndd: a network-based method for predicting drug-drug interactions. Nature Methods, 9(3):302–304, 2012.

[6] Hong Yu, Yilin Zhao, Yafang Li, and Maolin Wang. Ddinmf: a web server for predicting drug-drug interactions based on non-negative matrix factorization. BMC Systems Biology, 12(1):1–8, 2018.

[7] Jae Yong Ryu, Hwi-U Kim, and Sang-Yoon Lee. Deepddi: deep learning-based prediction of drug-drug interactions. Proceedings of the National Academy of Sciences, 115(18):E4304–E4311, 2018.

[8] Wei Lan, Jing Long, Jiawei Luo, Jiacheng He, Weiling Zeng, and Hui Liu. Ssi-ddi: substructure-substructure interaction-based deep neural network for drug–drug interaction prediction. Briefings in Bioinformatics, 22(5):bbab133, 2021.

[9] Qidong Liu, Yixiang Xie, Jiansheng Li, Yuhua Lu, and Jian-Xin Wang. Dsn-ddi: a dual-view substructure-aware graph neural network for drug–drug interaction prediction. Briefings in Bioinformatics, 24(1):bbac597, 2022.

[10] Guangyi Lin, Yuchen Zhang, Yijia Lin, Bo Xu, and Jingbo Zhou. A knowledge graph enhanced drug-drug interaction extraction approach for biomedical literature. Briefings in Bioinformatics, 24(1):bbac597, 2023.

[11] Jae-Hoon Lim, Sung-Hoon Ryu, Sung-Wook Kim, Jeong-Ah Heo, Hyun-Ju Lee, and Kwang-Seok Park. Dpddi: a deep predictor for drug-drug interactions. BMC Bioinformatics, 21(1):1–15, 2020.

[12] Xiaoping Lin, Ke Zhao, and Xiong Liu. Deeplgf: A new deep learning framework for drugdrug interaction prediction based on local and global feature fusion. Briefings in Bioinformatics, 23(6):bbac363, 2022.

[13] Yuancheng Liu, Yafang Li, and Maolin Wang. Gcnmk: a graph convolutional network with multikernel for drug-drug interaction prediction. Briefings in Bioinformatics, 23(2):bbab511, 2022.

[14] Haoran Wang, Jingtong Wang, Teng Zhang, and Yuedong Zhang. Graphsage-based drug-drug interaction prediction. Mathematical Biosciences and Engineering, 21(4):5559–5576, 2024.

[15] Yumo Yan, Jiaxin Huang, Yue Wu, and Feng Zhou. Drgatan: Dual-relational graph attention network for drug-drug interaction prediction. iScience, 27(7):109943, 2024.

[16] Ozan Kuru and A. Ercument Cicek. Mhcaddi: A multi-head co-attention based model for drug-drug interaction prediction. arXiv preprint 1905.00534, 2019.

[17] Shaked Brody, Uri Alon, and Eran Yahav. How attentive are graph attention networks? arXiv preprint 2105.14491, 2021.

[18] Ruochi Liu, Cheng Li, and Jing Wang. Gognn: A graph-of-graphs neural network for predicting protein-protein interactions. arXiv preprint 2005.05537, 2020.

[19] Zhaohui Deng, Yafang Li, and Maolin Wang. Dmfddi: a deep matrix factorization framework for drug-drug interaction prediction. Briefings in Bioinformatics, 24(6):bbad397, 2023.

[20] Zhenqin Zhu, Chao Shi, Yue Zhao, Daqing Liu, Shu Xu, Jiaxin Chen, Sheng Yu, and Mengjun Liu. Knowledge-guided pre-training of graph transformer for few-shot molecular property prediction. Nature Communications, 14(1):6566, 2023.

[21] Barun Bhhatarai, W. Patrick Walters, Cornelis E.C.A. Hop, Graeme Lanza, and Sean Ekins. Opportunities and challenges using artificial intelligence in adme/tox. Nature Materials, 18(5):418–422, 2019.

[22] Yujie Zhao, Yuchi Zhang, Zihan Li, Zelin Wang, Yinnian Liu, Yatao Liu, and Zaiqing Zhang. Moltc: A language-based task-agnostic molecular representation for drug discovery. arXiv preprint 2402.03781, 2024.

[23] Pan Zhou, Ziyang Wang, Yanjun Li, and Jia Li. Unigen-ddi: A unified generative framework for drug-drug interaction prediction. bioRxiv, 2024.

[24] Yutong Zhang, Zekun Liu, Jiawei Wang, and Jing Li. Ddi-ssl: A self-supervised learning framework for drug–drug interaction prediction. Applied Sciences, 13(19):10750, 2023.

[25] Yihang Chen, Yafang Li, and Maolin Wang. Deepgcl: A graph contrastive learning framework for drug-drug interaction prediction. PLoS ONE, 19(6):e0304798, 2024.

[26] Maxim Topaz, Diane L. Seger, and Sarah P. Slight. Alert fatigue in clinical decision support: reviewing the science and recommending a path forward. Journal of the American Medical Informatics Association, 23(6):1195–1200, 2016.

[27] Neda Rohani and Changiz Eslahchi. Mrhgnn: a multi-relational heterogeneous graph neural network for drug-drug interaction. Scientific Reports, 13(1):9988, 2023.

[28] Kexin Huang, Cao Xiao, Lucas M. Glass, and Jimeng Sun. Caster: Predicting drug interactions with chemical substructure representation. Proceedings of the AAAI Conference on Artificial Intelligence, 34(1):702–709, 2020.

[29] Qimai Li, Zhichao Han, and Xiao-Ming Wu. Deeper insights into graph convolutional networks for semi-supervised learning. In Thirty-Second AAAI Conference on Artificial Intelligence, 2018.

[30] Bibekananda Hati and Yogeshwar Singh Dadwhal. Pytorchlabflow: A local-first framework for reproducible deep learning experiments. Journal of Applied Bioanalysis, 11(S6):538–546, 2025.

[31] Yuxin Jiang, Yafang Li, and Maolin Wang. Drugman: a multi-level attention network for drug-drug interaction prediction. BMC Bioinformatics, 25(1):1–15, 2024.

